# Spatial attention and lateralized pre-stimulus alpha shape iconic memory via inhibition

**DOI:** 10.64898/2026.07.31.742006

**Authors:** Paul Justin Connor Smith, Niko A. Busch

## Abstract

The earliest stage of visual memory is often characterized as pre-attentive, yet recent work suggests that spatial attention and pre-stimulus alpha oscillations both shape iconic memory at its onset. Whether these influences reflect a common inhibitory mechanism or contribute independently is unknown. We combined an endogenous spatial pre-cue (valid, invalid, neutral) with an analysis of pre-stimulus alpha power in a partial-report paradigm, at target-to-post-cue onset asynchronies (SOAs) of 0, 120, and 1240 ms, while recording EEG from 45 participants. Behaviorally, perceptual sensitivity (*d^′^*) showed a cost-dominated signature: invalid cues impaired performance relative to both neutral and valid cues at the 0 and 120 ms SOAs, with no benefit of valid over neutral cues. The pre-stimulus EEG mirrored this asymmetry: at 120 ms, stronger alpha power ipsilateral to the target predicted higher *d^′^*, whereas contralateral alpha had no effect at any SOA, indicating suppression of the irrelevant hemifield rather than facilitation of the target. This ipsilateral effect was additive with cue validity rather than interactive, and it persisted in neutral trials without any directional cue. Iconic memory is thus continuously biased by the inhibitory state of early visual cortex, to which endogenous attention and spontaneous excitability fluctuations contribute through separable channels.

**Significance Statement:** Whether the earliest stage of visual memory depends on attention has long been debated. We show that the same asymmetry with which spatial attention shapes iconic memory—misdirecting attention impairs memory, but correctly directing it yields no benefit—also appears in EEG alpha power. Stronger prestimulus alpha power over the hemisphere ipsilateral to the target improves memory, while activity over the target-processing hemisphere does not. This alpha effect occurs even without any spatial cue, showing that it does not depend on deliberate attention. Because the cue-driven and spontaneous alpha effects add together rather than interact, they reflect two separable routes to the same outcome: suppression of irrelevant input. Early visual memory is thus continuously gated by the inhibitory state of visual cortex.

## Introduction

Attention is a fundamental cognitive process that selects relevant information while suppressing the irrelevant (Carrasco, 2011; Chun et al., 2011). While well established in high-level cognition (Hannula, 2018; P. L. Smith & Ratcliff, 2009), evidence for attentional modulation at earlier stages remains mixed, including in primary visual cortex (Martinez et al., 1999). This debate extends to iconic memory—a high-capacity, short-lived store characterized as either attention-independent or attention-modulated.

Iconic memory holds information from a brief glimpse available for selection and report, and is typically measured with partial-report paradigms: a multi-item array is followed by a post-cue at variable stimulus onset asynchronies (SOAs) indicating which item to report (Dick, 1974; Sperling, 1960). Accuracy is near-ceiling when array and post-cue coincide, then declines toward an asymptote over a few hundred milliseconds as information decays.

Influential models have argued that iconic memory is categorically pre-attentive, rooted in the distinction between phenomenal consciousness—the subjective experience of perceptual content—and access consciousness—information selected and made available for report and voluntary control of behavior. Iconic memory is placed on the phenomenal side: it holds rich contents that are consciously experienced yet not selected for access, with attention entering only at the transfer to short-term memory or an intermediate fragile store (Block, 1995; Lamme, 2004; Pinto et al., 2013). This view has been challenged by studies showing that diverting attention during encoding severely degrades iconic representations, sometimes to the point of inattentional blindness (Mack et al., 2015, 2016), though these effects may be post-perceptual (Aru & Bachmann, 2017).

Recent work located an attentional effect specifically at the onset of iconic memory (P. J. C. Smith & Busch, 2025a, 2025b, 2026). Using a spatial pre-cue at a 120 ms SOA, we found an asymmetry: misdirecting attention impaired accuracy (a cost), but correct direction yielded no benefit over baseline (P. J. C. Smith & Busch, 2026)—suppression of irrelevant information rather than enhancement of relevant information, arguing against a strictly pre-attentive account. Converging evidence comes from pre-stimulus brain states: both pupil-linked arousal and alpha power modulate iconic memory performance (P. J. C. Smith & Busch, 2025a, 2025b).

Alpha power (8–12 Hz) indexes neuronal excitability, with weaker power reflecting higher excitability (Buzsaki & Draguhn, 2004; Iemi et al., 2017). Iconic memory depends on persistent firing in early visual cortex (Teeuwen et al., 2021), whose gain is inversely related to alpha power (Dougherty et al., 2017); spatially specific alpha could therefore determine, hemifield by hemifield, which information persists. Per the gating-by-inhibition framework, alpha inhibits task-irrelevant cortical regions (Foxe & Snyder, 2011; Jensen & Mazaheri, 2010). Shifts of attention are tracked by alpha lateralization, with power increasing ipsilateral and decreasing contralateral to the attended location (Thut et al., 2006; Worden et al., 2000). Consistent with this, spontaneous trial-by-trial alpha lateralization extends the persistence of iconic memory via suppression at irrelevant locations, with no facilitatory contralateral effect (P. J. C. Smith & Busch, 2025b)—mirroring the cost of an invalid cue (P. J. C. Smith & Busch, 2026). Spontaneous alpha lateralization also predicts self-initiated shifts of spatial attention (Balestrieri & Busch, 2022; Bengson et al., 2014; Nadra et al., 2023), so spontaneous and cue-induced lateralization may reflect a common mechanism: if so, ipsilateral alpha power should index the same protective state regardless of its origin. We addressed this by combining a partial-report paradigm with endogenous spatial pre-cues (valid, invalid, neutral) and a post-cue at a target-to-post-cue SOA of 0, 120, or 1240 ms (Figure 1A), observing both cue-induced and spontaneous (neutral-trial) alpha lateralization and dissociating attentional costs from benefits.

**Figure 1:**
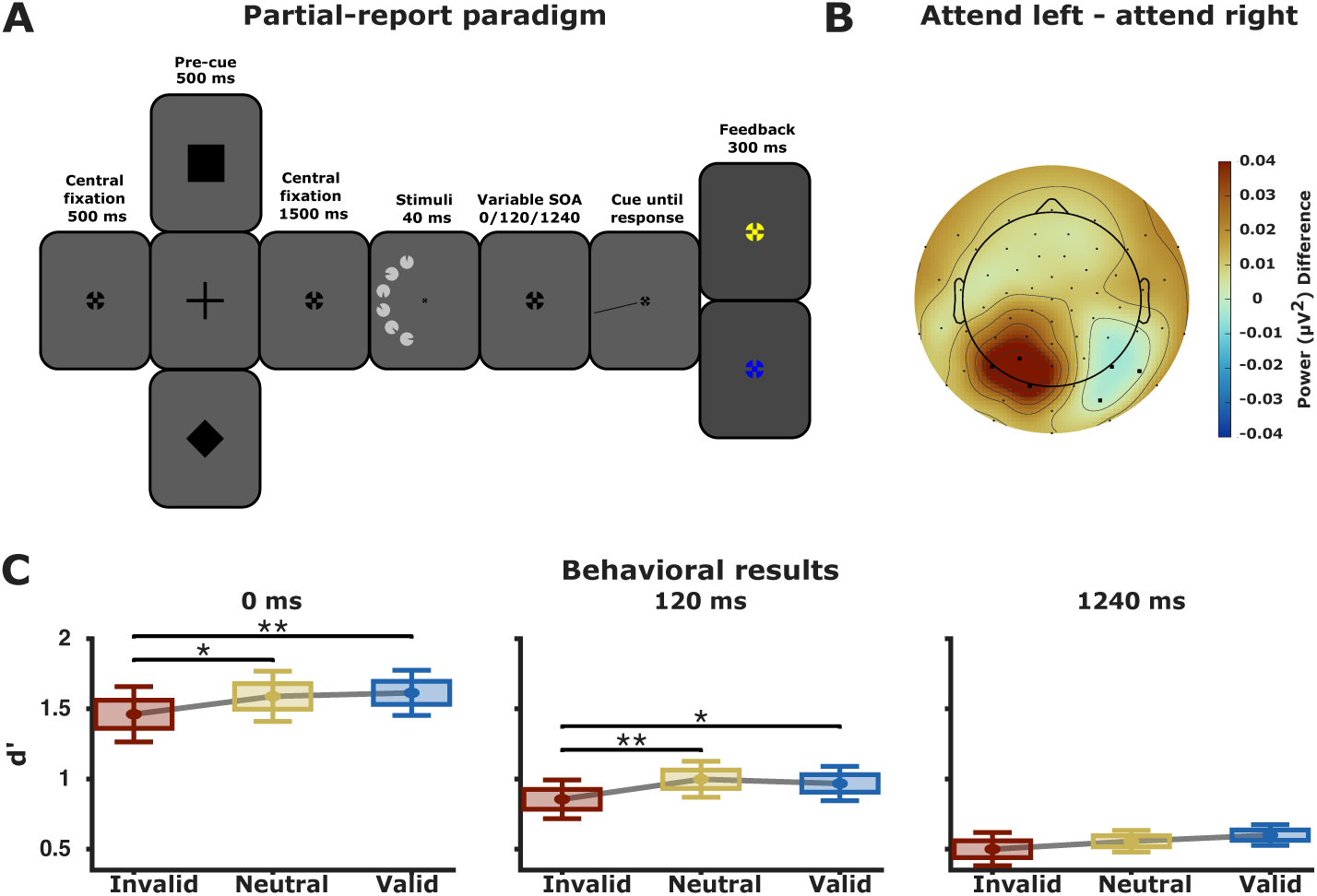
**A**: Illustration of the partial-report paradigm. Each trial began with a central fixation cross displayed for 500 ms. Following fixation, a pre-cue, which could either be a rectangle (indicating a stimulus presentation on the right side), a cross (indicating no cueing information), or a diamond (indicating a stimulus presentation on the left side) was presented for 500 ms. This pre-cue could either be valid (congruent with the subsequent target location), neutral or invalid (incongruent with the subsequent target location). A central fixation cross was again presented for 1500 ms. Following this, a stimulus array consisting of six items was presented on one side of the screen for 40 ms. Participants were then cued to report the orientation of one target item, with the post-cue appearing at various SOAs relative to target onset: at target onset, or 120 or 1240 ms after. There was no time limit for reporting or the subsequent confidence rating. After the confidence rating, feedback was given via the color of the fixation cross, turning blue for a correct response and yellow for an incorrect one. Image proportions are adjusted for illustration purposes. **B**: Topography plot showing the difference in pre-stimulus alpha power (8–12 Hz, 1000–2 ms before stimulus onset) between left cued and right cued spatial attention (both invalid and valid cues combined). Black electrodes indicate the binning electrodes. **C**: Accuracy (*d^′^*) data for invalid, neutral, and valid pre-cues at each target-to-post-cue SOA. Means of the conditions are connected with grey lines. Significant differences between the conditions are indicated with asterisks (* = *p <* 0.05; ** = *p <* 0.01).

We tested two pre-registered hypotheses: that invalid pre-cues would impair performance relative to neutral, with no benefit of valid over neutral, at 120 ms (P. J. C. Smith & Busch, 2026); and that stronger ipsilateral alpha power would predict better performance, with no effect of contralateral alpha (P. J. C. Smith & Busch, 2025b). Crucially, we tested whether pre-stimulus alpha and spatial attention reflect a shared inhibitory mechanism or independent contributions.

## Materials and Methods

### Pre-registration

The study was pre-registered with the Open Science Framework (https://osf.io/b95zr), and we adhered to the outlined methods unless stated otherwise.

### Participants

We collected behavioral, EEG, ECG, respiratory, and eye-tracking data from 55 healthy participants (aged 22.58 *±* 2.53 years, 42 female, 13 male). All participants had normal or corrected-to-normal vision, no reported history of neurological or psychiatric disorders, no reported history of heart diseases, provided their written consent, and were compensated with either course credits or money. The ethics commission of the faculty of psychology and sports sciences, University of Münster approved the study (ref. 2025-07-PS). Eight participants were excluded who misunderstood the assignment of cue symbols to attention conditions. Another two participants were excluded for a post-preprocessing trial count below 20%. The final sample size for the main analysis was 45 (aged 22.64 *±* 2.64 years), with 36 female and 9 male participants. A subset of the behavioral data of 40 of these subjects was already analyzed in P. J. C. Smith and Busch (2026).

### Stimuli and procedure

The experiment was presented on a 24-inch Viewpixx/EEG LCD Monitor with a 120 Hz refresh rate, 1 ms pixel response time, 95% luminance uniformity, and 1920×1080 pixels resolution (33.76×19.38; www.vpixx.com). The recording took place in a dimly lit, soundproof cabin. Participants’ heads were stabilized on a chinrest with their eyes approximately 86 cm from the monitor. Eye movements were monitored using a desktop-mounted Eyelink 1000+ infrared-based eye-tracking system (SR Research Ltd.) set to a 1000 Hz sampling rate (monocular, from the participant’s dominant eye).

The stimuli used in the partial-report paradigm were circles with wedges cut out at variable orientations (0*^◦^*, 45*^◦^*, 90*^◦^*, 135*^◦^*, 180*^◦^*, 225*^◦^*, 270*^◦^*, or 315*^◦^*). The circles had a diameter of 2.6 degrees visual angle (dva) whilst the cut-out had a thickness of 0.28 dva with a diameter of 0.6 dva. The circles were light grey (RGB: [180 180 180]), and the cut-out wedges were of a darker grey (RGB: [70 70 70]), matching the background color. The pre-cue utilized was a rectangle (RGB: [0 0 0]) for right-cued trials, a diamond (RGB: [0 0 0]) for left-cued trials, and a cross (RGB: [0 0 0]) for neutral trials. All pre-cues had the same dimensions as the stimuli (2.6 dva). The post-cue utilized was a black line (RGB: [0 0 0]) with a length of 1 dva and a thickness of 0.1 dva, pointing to one of the former positions of the stimuli.

Participants were instructed to avoid eye movements and blinks during the stimulus presentation. Participants had to fixate on the center of the screen, where a black (RGB: [0 0 0]) fixation cross with a size of 0.6 dva was presented for a fixed time interval of 500 ms. The pre-cue was presented for 500 ms after which the fixation cross was again presented for 1500 ms. Following this interval, six stimuli were arranged in a half circle to the left or to the right of the fixation cross and presented for 40 ms. The post-cue appeared after a variable SOA either at stimulus onset, or 120 or 1240 ms after stimulus onset. Participants were instructed to press a number on the number pad corresponding to the orientation of the wedge in the target circle (eight for 0*^◦^*, six for 90*^◦^*, two for 180*^◦^*, etc.). Following this decision, the participants were instructed to press either four (low), five (medium), or six (high) on the number pad, indicating their respective confidence rating. There was no time limit for their response. Feedback on their decision was provided 300 ms after their confidence rating, with correct responses marked by a blue (RGB: [0 0 255]) fixation cross, and incorrect responses indicated by a yellow (RGB: [255 255 0]) fixation cross. A visualization of a trial can be seen in Figure 1A. There were 200 training trials, where no data was collected. Following these training trials, there were 1000 trials, with short self-paced breaks after 200 consecutive trials. Post-cue SOA, stimulus position regarding the fixation cross, and target cut-out orientation were counterbalanced. There were a total of 175 invalid, 300 neutral, and 525 valid trials. The experiment was written and presented using Matlab2022 (mathworks.com), and Psychtoolbox (Brainard & Vision, 1997; Kleiner et al., 2007; Pelli, 1997).

### EEG acquisition and pre-processing

EEG activity was recorded using a Biosemi Active Two EEG system with 67 Ag/AgCl electrodes (BioSemi B.V.) set to a 1024 Hz sampling rate. Sixty-four electrodes were arranged in a custom-made montage with equidistant placement (EASYCAP GmbH) with additional external electrodes placed next to the right eye, the left eye, and below the left eye.

All EEG data preprocessing and analysis steps were scripted and run in Matlab2025a (mathworks.co m) using the EEGLAB 2023.1 toolbox (Delorme & Makeig, 2004) and custom scripts. The continuous data was downsampled to 256 Hz and re-referenced to the average. It was then high-pass filtered at 0.1 Hz, and low-pass filtered at 40 Hz. Following this, the data was epoched from −1500 ms to 1500 ms time-locked to stimulus onset. A round of trial rejection based on thresholding (*±* 500 µV) and joint probability (function pop_jointprob in EEGLAB, local threshold 9, global threshold 5) was performed. Furthermore, trials in which eyeblinks were detected in a range of −500 ms to 500 ms around stimulus onset, or in which participants significantly deviated from the fixation cross (2.5 dva) were rejected. An average of 296.3 trials were excluded per participant. After this, ICA was applied, and components were identified by the IClabel algorithm (Pion-Tonachini et al., 2019) as “Brain”, “Muscle”, “Eye”, “Heart”, “Line Noise”, “Channel Noise”, or “Other”. Components were manually screened, and components classified as non-brain activity were excluded. Following this, noisy channels (SD *>* 2) were spherically interpolated. Three participants each had one channel interpolated.

### EEG data analysis

Pre-stimulus spectral EEG power was computed using a Fast Fourier Transform (FFT) of the data within the pre-stimulus time range from −1000 ms to −2 ms before stimulus onset for all electrodes and frequencies. The time window deviated from the preregistration because cueing-related alpha modulation extended beyond the preregistered time window of −500 to −2 ms (see e.g. Figure 3). Single-trial prestimulus alpha power in the frequency range of 8–12 Hz was extracted at electrode locations P3, P5, P6, P8, PO3, and PO8, selected on the basis of visual inspection of the topographic distribution of attention-related alpha modulation (see Figure 1B). Ipsi- and contralateral pre-stimulus alpha power relative to the visual hemifield of the stimuli presentation was extracted. Left (P3, P5, PO3) and right (P6, P8, PO8) electrodes were chosen based on the electrodes of the bilateral analysis. A pre-stimulus lateralization index was computed by subtracting contralateral pre-stimulus alpha power relative to the target from ipsilateral pre-stimulus alpha power relative to the target and normalizing this by dividing it through the sum of both: 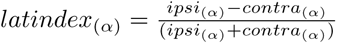 (Thut et al., 2006). Negative values for this lateralization index indicate relatively more contralateral pre-stimulus alpha power while positive values indicate relatively more ipsilateral pre-stimulus alpha power.

Alpha power and behavioral performance are both susceptible to time-on-task effects, meaning that each can drift systematically over the course of an experiment due to fatigue, practice, or gradual changes in alertness and cognitive strategy (Benwell et al., 2019; Kopčanová et al., 2024). If left uncontrolled, such drifts can induce a spurious correlation between alpha power and performance, since both measures may co-vary with time rather than with one another. To address this, each single-trial alpha power and lateralization estimate was ranked and binned into tertiles using a sliding window approach. For each trial, the local distribution of alpha power was estimated from a window of 40 adjacent trials (corresponding to a time span of *≈* 3 minutes) centered on that trial, and the trial’s bin assignment reflected its rank within this local distribution rather than its rank across the full session. This procedure removed slow monotonic trends from the binning while preserving trial-to-trial variability in alpha power.

For each pre-cue condition and each target-to-post-cue SOA, *d^′^* was computed across all pre-stimulus alpha power measure bins (J. K. Smith, 1982).

### Statistical analysis

To replicate the effect of pre-cue validity on *d^′^* found in P. J. C. Smith and Busch (2026), *d^′^*at each target-to-post-cue SOA was assessed using a single-factor linear mixed-effects model (LME) with pre-cue validity (invalid, neutral, valid) as a fixed effect and random intercepts per subject:

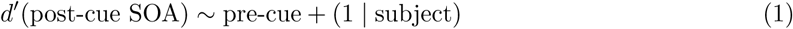

To assess the joint and interactive effects of pre-cue validity and alpha power, each alpha predictor was then entered into a two-factor interaction model together with pre-cue validity:

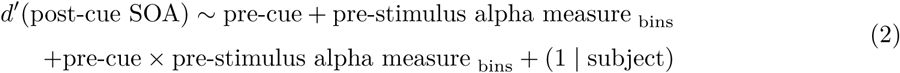

where pre-stimulus alpha measure _bins_ was one of four predictors entered separately: bilateral prestimulus alpha power, pre-stimulus alpha lateralization, ipsilateral pre-stimulus alpha power, or contralateral pre-stimulus alpha power relative to the target.

Pairwise post-hoc Wald *F*-tests were computed for all significant effects by applying contrast vectors to the fitted LME model. Cohen’s *d* was computed as the contrast estimate divided by the residual standard deviation of the model, providing a within-subject standardized effect size.

In deviation from the pre-registration, we decided against testing separate single-factor models for the pre-stimulus alpha predictors as the two-factor models allowed us to better analyze the joint effects and interactions between the spatial attention manipulation and pre-stimulus alpha. To test for the effect found in P. J. C. Smith and Busch (2025b) we decided to analyze whether pre-stimulus alpha power modulates perceptual sensitivity independently of directed spatial attention. All four alpha power predictors were tested within neutral pre-cue trials only, using one-way LMEs of the form of

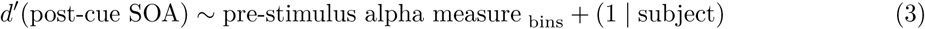

fit to the neutral trial subset. This allowed for a more accurate replication compared to the intended single-factor models as the neutral subset is more similar to the experimental setup of the partial-report paradigm in P. J. C. Smith and Busch (2025b).

To investigate any potential influences of aperiodic activity of our data on the analysis results, we applied specparam (Donoghue et al., 2020). We compared the aperiodic exponent and offset for the averaged correct and incorrect trials at our binning electrodes using *t*-tests.

As alpha lateralization has been associated with eye movements and miniature gaze shifts (Mössing et al., 2024; Popov et al., 2023; Van Ede et al., 2019), we investigated the spatial distribution of gaze shifts and their relationship with performance. We computed the two-dimensional histogram of gaze positions in the same time interval used for the analysis of pre-stimulus alpha power (−1000 to −2 ms), separately for correct and incorrect trials, for each pre-cueing condition and for left- and right-hemifield targets at the 120 ms post-cue SOA. These histograms were then convolved with a 2D Gaussian (sigma = 15 pixels) to compute gaze density, represented as a heat map. We then tested for differences between gaze density on correct and incorrect trials at each pixel using two-sided *t*-tests, correcting for multiple comparisons across pixels using the false discovery rate (FDR; *q <* 0.05).

### Data and code accessibility

The data will be available upon acceptance of the manuscript at the Open Science Framework (https://osf.io/b95zr). The code will be available at https://github.com/pauljcs/alpha-attention-iconic.

## Results

### Behavioral results

A significant main effect of pre-cue validity on *d^′^* was observed at the 0 ms target-to-post-cue SOA (*F* (2, 132) = 3.97, *p* = 0.021) and at the 120 ms target-to-post-cue SOA (*F* (2, 132) = 5.14, *p* = 0.007), but not at the 1240 ms target-to-post-cue SOA (*F* (2, 132) = 1.81, *p* = 0.168). Post-hoc tests at 0 ms revealed that *d^′^* was significantly lower for invalid than for neutral pre-cues (*F* (1, 132) = 4.83, *p* = 0.030, *d* = 0.46) as well as for invalid in comparison to valid pre-cues (*F* (1, 132) = 6.90, *p* = 0.010, *d* = 0.55). No significant difference between neutral and valid pre-cues was observed (*F* (1, 132) = 0.18, *p* = 0.668). Post-hoc tests at 120 ms revealed that *d^′^* was significantly lower for invalid than for neutral pre-cues (*F* (1, 132) = 9.26, *p* = 0.003, *d* = 0.64) and lower for invalid than for valid pre-cues (*F* (1, 132) = 5.74, *p* = 0.018, *d* = 0.51). Neutral and valid pre-cues did not differ (*F* (1, 132) = 0.42, *p* = 0.519; see Figure 1C).

These results largely replicate the results from P. J. C. Smith and Busch (2026), which is expected as they share some subject data. The significant effect of invalid pre-cues being associated with lower *d^′^*in comparison with valid pre-cues at the 1240 ms target-to-post-cue SOA in P. J. C. Smith and Busch (2026) could not be observed. A visualization of the difference in pre-stimulus alpha power between left cued and right cued spatial attention showed a clear pattern of alpha lateralization (see Figure 1B).

### Pre-stimulus alpha and iconic memory performance

We next tested whether pre-stimulus alpha power modulated these behavioral effects, entering each alpha measure into a two-factor model with pre-cue validity.

#### Bilateral pre-stimulus alpha

A significant main effect of pre-cue validity was observed at the 120 ms target-to-post-cue SOA (*F* (2, 395) = 5.01, *p* = 0.007), with no significant main effect of bilateral pre-stimulus alpha power (*F* (2, 395) = 2.19, *p* = 0.114) or interaction (*F* (4, 395) = 0.71, *p* = 0.585; see Figure 2A). Post-hoc tests revealed the same pattern as the single factor pre-cue model above.

**Figure 2:**
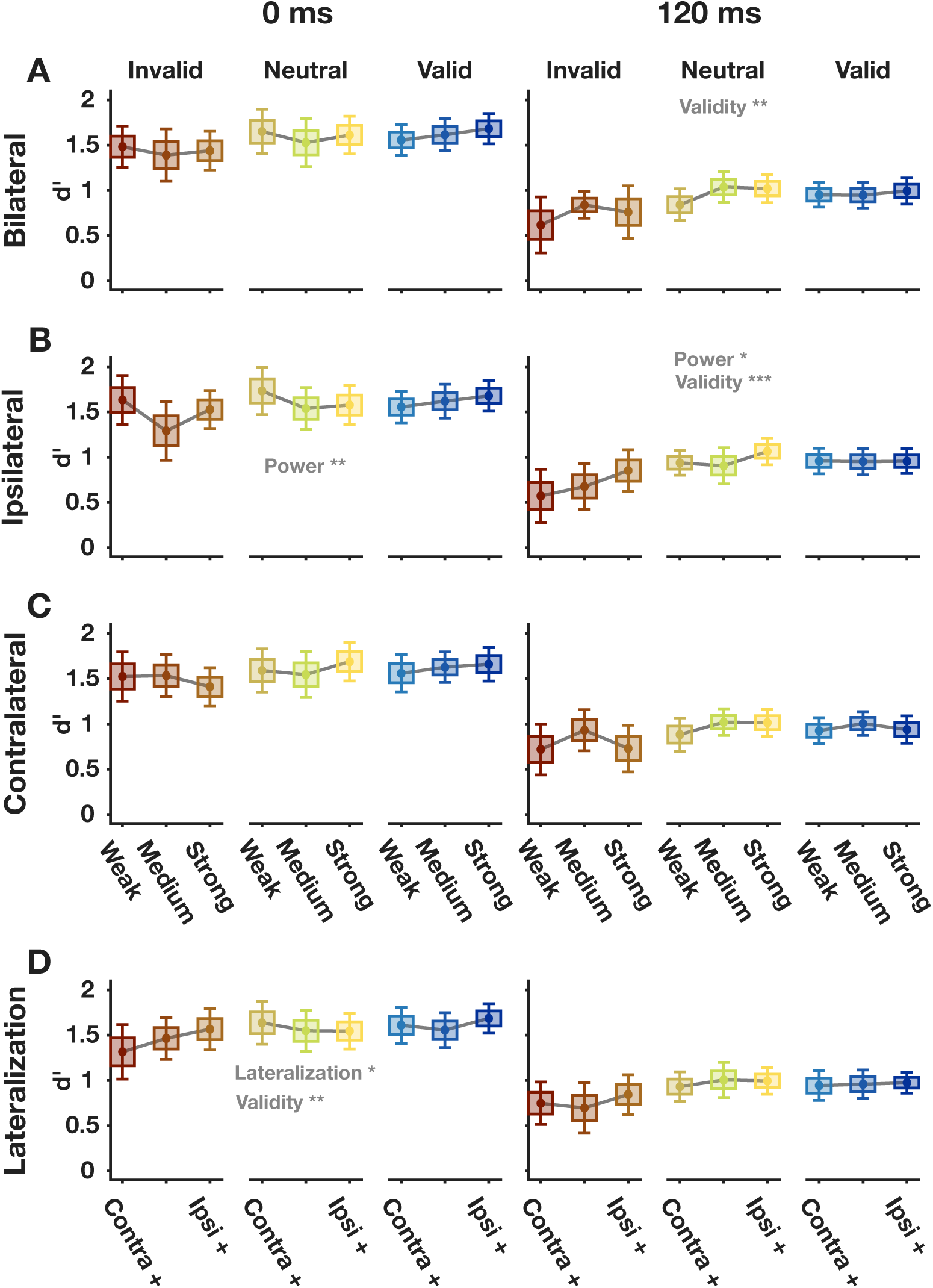
**A**: Accuracy (*d^′^*) data for weak, medium and strong bilateral pre-stimulus alpha power bins for invalid, neutral and valid trials at the 0 and 120 ms target-to-post-cue SOA. Conditions data is shown separately for invalid, neutral and valid trials. Means of the conditions are connected with grey lines. Significant main effects across the models are indicated with grey text and asterisks (* = *p <* 0.05; ** = *p <* 0.01). All of the following panels follow these conventions. **B**: Accuracy (*d^′^*) data for weak, medium and strong ipsilateral pre-stimulus alpha power relative to the target bins for invalid, neutral and valid trials at the 0 and 120 ms target-to-post-cue SOA. **C**: Accuracy (*d^′^*) data for weak, medium and strong contralateral pre-stimulus alpha power relative to the target bins for invalid, neutral and valid trials at the 0 and 120 ms target-to-post-cue SOA. **D**: Accuracy (*d^′^*) data for weak, medium and strong pre-stimulus alpha lateralization relative to the target for invalid, neutral and valid trials bins at the 0 and 120 ms target-to-post-cue SOA.

No significant effects were observed at either the 0 or 1240 ms target-to-post-cue SOAs (0: all *F ≤* 1.5, all *p ≥* 0.225; 1240: all *p ≥* 0.114, all *F ≤* 2.18). The full results are reported in Table 1.

**Table 1:**
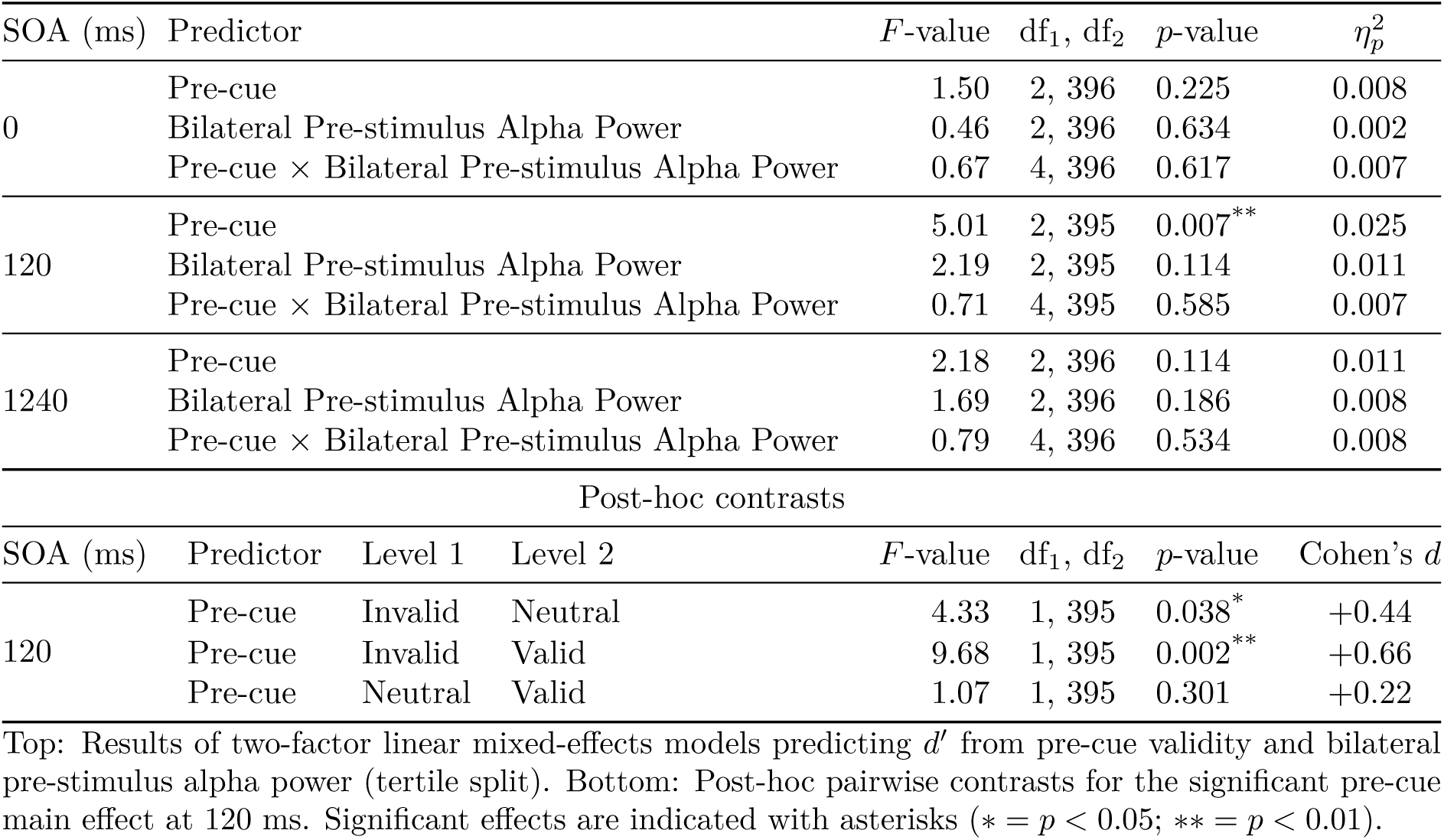
Bilateral pre-stimulus alpha.

| SOA (ms) | Predictor | | $F$-value | df<sub>1</sub>, df<sub>2</sub> | $p$-value | $\eta_p^2$ | |
| --- | --- | --- | --- | --- | --- | --- | --- |
| 0 | Pre-cue |  | 1.50 | 2, 396 | 0.225 | 0.008 |  |
|  | Bilateral Pre-stimulus Alpha Power |  | 0.46 | 2, 396 | 0.634 | 0.002 |  |
| | Pre-cue × Bilateral Pre-stimulus Alpha Power | | 0.67 | 4, 396 | 0.617 | 0.007 | |
| 120 | Pre-cue |  | 5.01 | 2, 395 | 0.007** | 0.025 |  |
|  | Bilateral Pre-stimulus Alpha Power |  | 2.19 | 2, 395 | 0.114 | 0.011 |  |
| | Pre-cue × Bilateral Pre-stimulus Alpha Power | | 0.71 | 4, 395 | 0.585 | 0.007 | |
| 1240 | Pre-cue |  | 2.18 | 2, 396 | 0.114 | 0.011 |  |
|  | Bilateral Pre-stimulus Alpha Power |  | 1.69 | 2, 396 | 0.186 | 0.008 |  |
| | Pre-cue × Bilateral Pre-stimulus Alpha Power | | 0.79 | 4, 396 | 0.534 | 0.008 | |
| Post-hoc contrasts |  |  |  |  |  |  |  |
| SOA (ms) | Predictor | Level 1 | Level 2 | $F$-value | df<sub>1</sub>, df<sub>2</sub> | $p$-value | Cohen's $d$ |
| 120 | Pre-cue | Invalid | Neutral | 4.33 | 1, 395 | 0.038* | +0.44 |
|  | Pre-cue | Invalid | Valid | 9.68 | 1, 395 | 0.002** | +0.66 |
|  | Pre-cue | Neutral | Valid | 1.07 | 1, 395 | 0.301 | +0.22 |

Top: Results of two-factor linear mixed-effects models predicting $d'$ from pre-cue validity and bilateral pre-stimulus alpha power (tertile split). Bottom: Post-hoc pairwise contrasts for the significant pre-cue main effect at 120 ms. Significant effects are indicated with asterisks ( $*$ = $p < 0.05$ ; $**$ = $p < 0.01$ ).

#### Pre-stimulus alpha lateralization

At the 0 ms target-to-post-cue SOA, significant main effects of both pre-cue validity (*F* (2, 396) = 6.33, *p* = 0.002) and pre-stimulus alpha lateralization relative to the target (*F* (2, 396) = 3.18, *p* = 0.043) were observed, with no significant interaction (*F* (4, 396) = 1.68, *p* = 0.153; see Figure 2D and Figure 3A, C & E). The pre-cue validity effect followed the established pattern of invalid *<* neutral and invalid *<* valid with neutral and valid not differing. Post-hoc tests of the lateralization main effect revealed a significant difference between weak and strong bins (*F* (1, 396) = 6.28, *p* = 0.013, *d* = 0.53) with strong lateralization resulting in better performance and no significant contrasts between weak and medium (*F* (1, 396) = 2.22, *p* = 0.137) or medium and strong (*F* (1, 396) = 1.03, *p* = 0.310; see Figure 2D).

**Figure 3:**
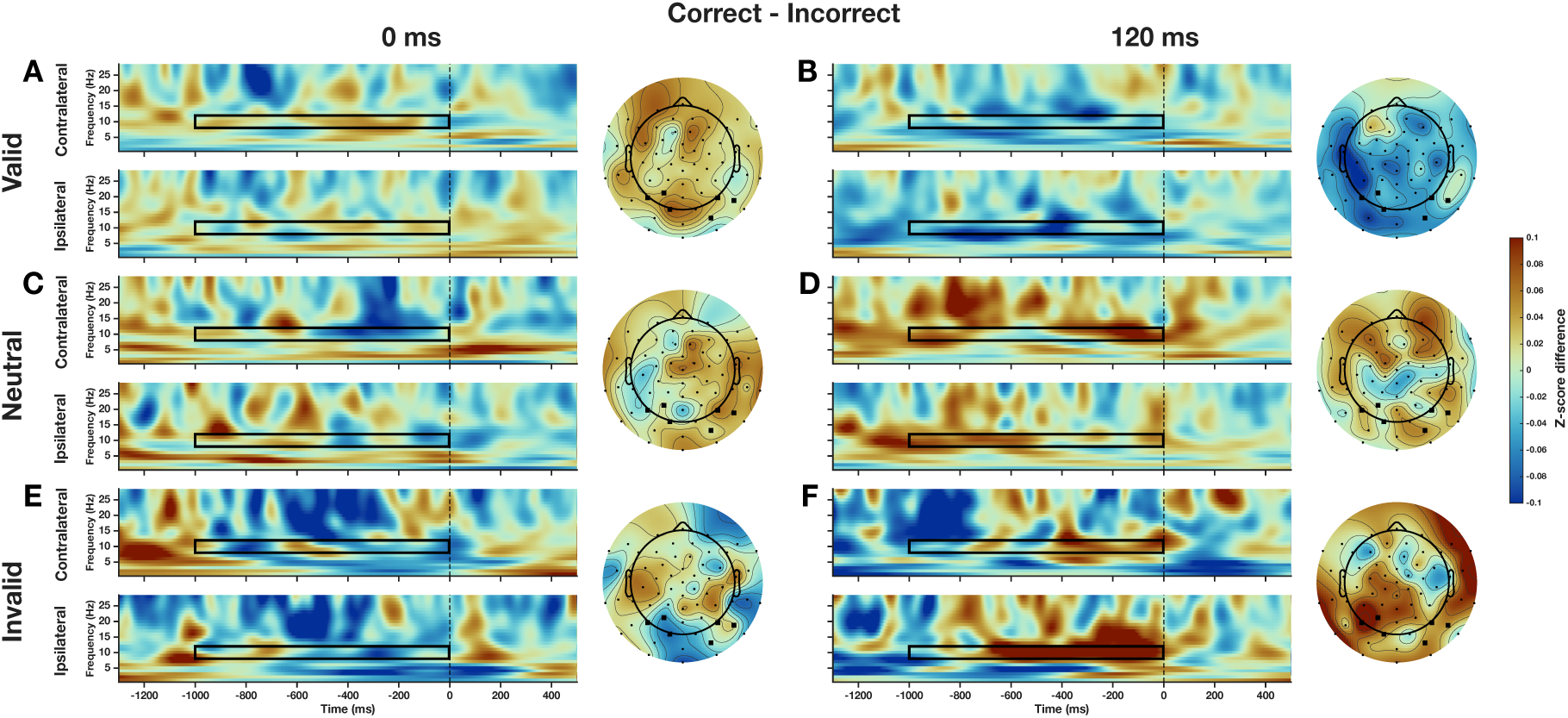
**A**: Time-frequency and topography plots showing the difference in z-scored pre-stimulus alpha power relative to the target at the 0 ms target-to-post-cue SOA for trials with a valid pre-cue (spatial attention cued to the hemisphere of the subsequent target). Upper left time-frequency plot shows the difference in z-scored power at the binning electrodes contralateral to the presented stimulus. Lower left time-frequency plot shows the difference in z-scored power at the binning electrodes ipsilateral to the presented stimulus. The black rectangle indicates the time and frequency window used in the binning. The dashed line indicates the time of the stimulus presentation. Right topography plot shows the z-scored pre-stimulus alpha power difference (8–12 Hz, 1000–2 ms before target onset) where right-target trials were hemisphere-flipped prior to averaging so that the left side of each scalp map consistently represents the ipsilateral hemisphere and the right side represents the contralateral hemisphere relative to the target. Black electrodes indicate the binning electrodes. All of the following panels follow these conventions. **B**: Time-frequency and topography plots showing the difference in z-scored pre-stimulus alpha power relative to the target at the 120 ms target-to-post-cue SOA for trials with a valid pre-cue. **C**: Time-frequency and topography plots showing the difference in z-scored pre-stimulus alpha power relative to the target at the 0 ms target-to-post-cue SOA for trials with a neutral pre-cue (no directional spatial attention cueing). **D**: Time-frequency and topography plots showing the difference in z-scored pre-stimulus alpha power relative to the target at the 120 ms target-to-post-cue SOA for trials with a neutral pre-cue. **E**: Time-frequency and topography plots showing the difference in z-scored pre-stimulus alpha power relative to the target at the 0 ms target-to-post-cue SOA for trials with an invalid pre-cue (spatial attention cued to the opposite hemisphere of the subsequent target presentation). **F**: Time-frequency and topography plots showing the difference in z-scored pre-stimulus alpha power relative to the target at the 120 ms target-to-post-cue SOA for trials with an invalid pre-cue.

No significant effects were observed at 120 ms (all *F ≤* 2.38, all *p ≥* 0.094; see Figure 2D).

At 1240 ms, a significant main effect of pre-stimulus alpha lateralization was observed (*F* (2, 396) = 3.46, *p* = 0.032), with no significant pre-cue effect (*F* (2, 396) = 0.07, *p* = 0.928) or interaction (*F* (4, 396) = 1.37, *p* = 0.245). Post-hoc contrasts revealed lower *d^′^* for medium compared to weak lateralization (*F* (1, 396) = 5.69, *p* = 0.018, *d* = *−*0.50) and for strong compared to weak lateralization (*F* (1, 396) = 4.64, *p* = 0.032, *d* = *−*0.45), with no significant difference between medium and strong (*F* (1, 396) = 0.05, *p* = 0.818). The full results are reported in Table 2.

**Table 2:** Pre-stimulus alpha lateralization.

| SOA (ms) | Predictor |  | *F*-value | df<sub>1</sub>, df<sub>2</sub> | *p*-value | η<sub>*p*</sub><sup>2</sup> |  |
| --- | --- | --- | --- | --- | --- | --- | --- |
| 0 | Pre-cue |  | 6.33 | 2, 396 | 0.002** | 0.031 |  |
|  | Pre-stimulus Alpha Lateralization |  | 3.18 | 2, 396 | 0.043* | 0.016 |  |
|  | Pre-cue × Pre-stimulus Alpha Lateralization |  | 1.68 | 4, 396 | 0.153 | 0.017 |  |
| 120 | Pre-cue |  | 2.38 | 2, 396 | 0.094 | 0.012 |  |
|  | Pre-stimulus Alpha Lateralization |  | 1.13 | 2, 396 | 0.325 | 0.006 |  |
|  | Pre-cue × Pre-stimulus Alpha Lateralization |  | 0.41 | 4, 396 | 0.802 | 0.004 |  |
| 1240 | Pre-cue |  | 0.07 | 2, 396 | 0.928 | 0.000 |  |
|  | Pre-stimulus Alpha Lateralization |  | 3.46 | 2, 396 | 0.032* | 0.017 |  |
|  | Pre-cue × Pre-stimulus Alpha Lateralization |  | 1.37 | 4, 396 | 0.245 | 0.014 |  |
| Post-hoc contrasts |  |  |  |  |  |  |  |
| SOA (ms) | Predictor | Level 1 | Level 2 | *F*-value | df<sub>1</sub>, df<sub>2</sub> | *p*-value | Cohen's *d* |
| 0 | Pre-cue | Invalid | Neutral | 10.29 | 1, 396 | 0.001** | +0.68 |
|  | Pre-cue | Invalid | Valid | 8.63 | 1, 396 | 0.003** | +0.62 |
|  | Pre-cue | Neutral | Valid | 0.07 | 1, 396 | 0.788 | −0.06 |
|  | Lateralization | Weak | Medium | 2.22 | 1, 396 | 0.137 | +0.31 |
|  | Lateralization | Weak | Strong | 6.28 | 1, 396 | 0.013* | +0.53 |
|  | Lateralization | Medium | Strong | 1.03 | 1, 396 | 0.310 | +0.21 |
| 1240 | Lateralization | Weak | Medium | 5.69 | 1, 396 | 0.018* | −0.50 |
|  | Lateralization | Weak | Strong | 4.64 | 1, 396 | 0.032* | −0.45 |
|  | Lateralization | Medium | Strong | 0.05 | 1, 396 | 0.818 | +0.05 |

Top: Results of two-factor linear mixed-effects models predicting $d'$ from pre-cue validity and pre-stimulus alpha lateralization relative to the target (tertile split). Bottom: Post-hoc pairwise contrasts for significant main effects. Significant effects are indicated with asterisks (\* = $p < 0.05$ ; \*\* = $p < 0.01$ ).

#### Ipsilateral pre-stimulus alpha

At the 0 ms target-to-post-cue SOA, a significant main effect of ipsilateral pre-stimulus alpha power relative to the target (*F* (2, 396) = 5.01, *p* = 0.007) and a trend for the interaction (*F* (4, 396) = 2.08, *p* = 0.083) were observed, with no pre-cue main effect (*F* (2, 396) = 1.29, *p* = 0.276; see Figure 2B and Figure 3A, C, & E). Post-hoc tests of the ipsilateral pre-stimulus alpha main effect revealed that *d^′^*was significantly lower for medium compared to weak ipsilateral pre-stimulus alpha power (*F* (1, 396) = 9.57, *p* = 0.002, *d* = *−*0.65) and significantly higher for strong compared to medium (*F* (1, 396) = 4.52, *p* = 0.034, *d* = 0.45), with no significant difference between weak and strong (*F* (1, 396) = 0.93, *p* = 0.335).

At 120 ms, significant main effects of both pre-cue validity (*F* (2, 396) = 8.45, *p* = 0.0003) and ipsilateral pre-stimulus alpha power relative to the target (*F* (2, 396) = 3.58, *p* = 0.029) were observed, with no interaction (*F* (4, 396) = 1.02, *p* = 0.397; see Figure 2B and Figure 3B, D, & F). The precue validity effect followed the established pattern. Post-hoc tests of the ipsilateral pre-stimulus alpha power effect revealed that strong power resulted in significantly better performance than weak power (*F* (1, 396) = 7.01, *p* = 0.008, *d* = 0.56), with no other contrasts reaching significance (all *p ≥* 0.097).

At 1240 ms, a significant pre-cue effect (*F* (2, 396) = 4.75, *p* = 0.009) and a trend for ipsilateral pre-stimulus alpha power relative to the target (*F* (2, 396) = 2.96, *p* = 0.053) were observed, with no interaction (*F* (4, 396) = 1.17, *p* = 0.325). The pre-cue validity effect followed the established pattern. The full results are reported in Table 3.

**Table 3:** Ipsilateral pre-stimulus alpha power.

| SOA (ms) | Predictor | | $F$-value | df<sub>1</sub>, df<sub>2</sub> | $p$-value | $\eta_p^2$ | |
| --- | --- | --- | --- | --- | --- | --- | --- |
| 0 | Pre-cue |  | 1.29 | 2, 396 | 0.276 | 0.006 |  |
|  | Ipsilateral Pre-stimulus Alpha Power |  | 5.01 | 2, 396 | 0.007** | 0.025 |  |
| | Pre-cue × Ipsilateral Pre-stimulus Alpha Power | | 2.08 | 4, 396 | 0.083 | 0.021 | |
| 120 | Pre-cue |  | 8.45 | 2, 396 | 0.0003** | 0.041 |  |
|  | Ipsilateral Pre-stimulus Alpha Power |  | 3.58 | 2, 396 | 0.029* | 0.018 |  |
| | Pre-cue × Ipsilateral Pre-stimulus Alpha Power | | 1.02 | 4, 396 | 0.397 | 0.010 | |
| 1240 | Pre-cue |  | 4.75 | 2, 396 | 0.009** | 0.023 |  |
|  | Ipsilateral Pre-stimulus Alpha Power |  | 2.96 | 2, 396 | 0.053 | 0.015 |  |
| | Pre-cue × Ipsilateral Pre-stimulus Alpha Power | | 1.17 | 4, 396 | 0.325 | 0.012 | |
| Post-hoc contrasts |  |  |  |  |  |  |  |
| SOA (ms) | Predictor | Level 1 | Level 2 | $F$-value | df<sub>1</sub>, df<sub>2</sub> | $p$-value | Cohen's $d$ |
| 0 | Ipsilateral Alpha | Weak | Medium | 9.57 | 1, 396 | 0.002** | −0.65 |
|  | Ipsilateral Alpha | Weak | Strong | 0.93 | 1, 396 | 0.335 | −0.20 |
|  | Ipsilateral Alpha | Medium | Strong | 4.52 | 1, 396 | 0.034* | +0.45 |
| 120 | Pre-cue | Invalid | Neutral | 11.97 | 1, 396 | 0.001** | +0.73 |
|  | Pre-cue | Invalid | Valid | 13.36 | 1, 396 | 0.0003** | +0.77 |
|  | Pre-cue | Neutral | Valid | 0.04 | 1, 396 | 0.845 | +0.04 |
|  | Ipsilateral Alpha | Weak | Medium | 0.97 | 1, 396 | 0.326 | +0.21 |
|  | Ipsilateral Alpha | Weak | Strong | 7.01 | 1, 396 | 0.008** | +0.56 |
|  | Ipsilateral Alpha | Medium | Strong | 2.77 | 1, 396 | 0.097 | +0.35 |
| 1240 | Pre-cue | Invalid | Neutral | 6.22 | 1, 396 | 0.013* | +0.53 |
|  | Pre-cue | Invalid | Valid | 7.94 | 1, 396 | 0.005** | +0.59 |
|  | Pre-cue | Neutral | Valid | 0.10 | 1, 396 | 0.746 | +0.07 |

Top: Results of two-factor linear mixed-effects models predicting $d'$ from pre-cue validity and ipsilateral pre-stimulus alpha power (tertile split). Bottom: Post-hoc pairwise contrasts for significant main effects. Significant effects are indicated with asterisks (\* = $p < 0.05$ ; \*\* = $p < 0.01$ ).

#### Contralateral pre-stimulus alpha

Finally, we examined contralateral pre-stimulus alpha power relative to the target. No significant main effects or interactions were observed at any target-to-post-cue SOA (see Figure 2C, Figure 3, and Table 4).

**Table 4:**
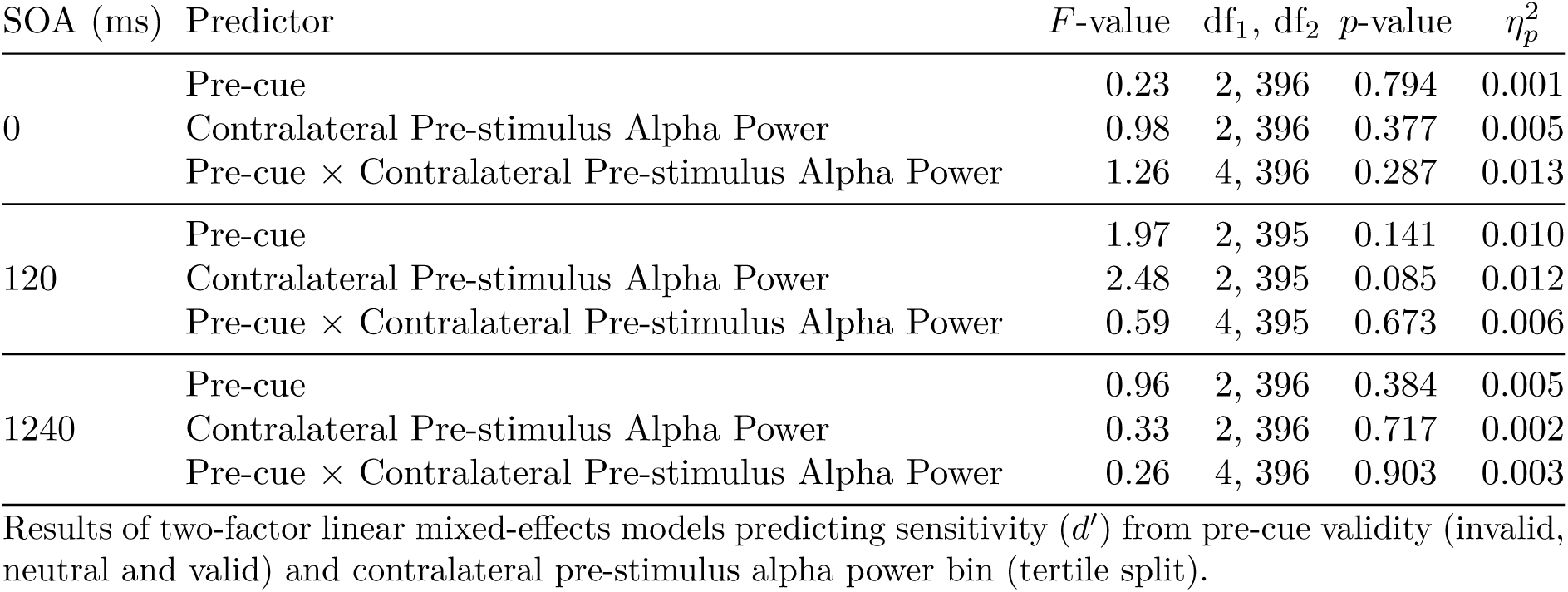
Contralateral pre-stimulus alpha power.

| SOA (ms) | Predictor | $F$-value | df<sub>1</sub>, df<sub>2</sub> | $p$-value | $\eta_p^2$ |
| --- | --- | --- | --- | --- | --- |
| 0 | Pre-cue | 0.23 | 2, 396 | 0.794 | 0.001 |
|  | Contralateral Pre-stimulus Alpha Power | 0.98 | 2, 396 | 0.377 | 0.005 |
| | Pre-cue × Contralateral Pre-stimulus Alpha Power | 1.26 | 4, 396 | 0.287 | 0.013 |
| 120 | Pre-cue | 1.97 | 2, 395 | 0.141 | 0.010 |
|  | Contralateral Pre-stimulus Alpha Power | 2.48 | 2, 395 | 0.085 | 0.012 |
| | Pre-cue × Contralateral Pre-stimulus Alpha Power | 0.59 | 4, 395 | 0.673 | 0.006 |
| 1240 | Pre-cue | 0.96 | 2, 396 | 0.384 | 0.005 |
|  | Contralateral Pre-stimulus Alpha Power | 0.33 | 2, 396 | 0.717 | 0.002 |
| | Pre-cue × Contralateral Pre-stimulus Alpha Power | 0.26 | 4, 396 | 0.903 | 0.003 |

Results of two-factor linear mixed-effects models predicting sensitivity ( $d'$ ) from pre-cue validity (invalid, neutral and valid) and contralateral pre-stimulus alpha power bin (tertile split).

### Pre-stimulus alpha and performance without directed attention

To test whether pre-stimulus alpha power modulated perceptual sensitivity independently of directed spatial attention, all four alpha power predictors were tested within neutral pre-cue trials only.

No significant effect of pre-stimulus alpha lateralization or contralateral pre-stimulus alpha power relative to the target was observed at any target-to-post-cue SOA (lateralization: all *F ≤* 1.89, all *p ≥* 0.154; contralateral: all *F ≤* 2.01, all *p ≥* 0.138).

A significant main effect of bilateral pre-stimulus alpha power was observed at 120 ms (*F* (2, 160) = 3.25, *p* = 0.042; see Figure 2A and Figure 3D). Post-hoc contrasts revealed lower *d^′^*for weak compared to medium bilateral pre-stimulus alpha power (*F* (1, 160) = 5.07, *p* = 0.026, *d* = 0.43) and lower *d^′^* for weak compared to strong bilateral pre-stimulus alpha power (*F* (1, 160) = 4.67, *p* = 0.032, *d* = 0.42), with no significant difference between medium and strong (*F* (1, 160) = 0.009, *p* = 0.925). A significant main effect of ipsilateral pre-stimulus alpha power relative to the target was also observed at 120 ms (*F* (2, 161) = 3.97, *p* = 0.021; see Figure 2B and Figure 3D). Post-hoc contrasts revealed significantly higher *d^′^* for strong compared to weak ipsilateral pre-stimulus alpha power (*F* (1, 161) = 4.51, *p* = 0.035, *d* = 0.41) and significantly higher *d^′^* for strong compared to medium ipsilateral pre-stimulus alpha power (*F* (1, 161) = 7.09, *p* = 0.009, *d* = 0.51), with no significant difference between weak and medium (*F* (1, 161) = 0.27, *p* = 0.601). No significant effects of bilateral or ipsilateral pre-stimulus alpha power were observed at the 0 ms or the 1240 ms target-to-post-cue SOA (bilateral: all *F ≤* 0.38, all *p ≥* 0.685; ipsilateral: all *F ≤* 2.02, all *p ≥* 0.136; see Table 5 and Figure 2).

**Table 5:** Alpha power effects on *d^′^* within neutral pre-cue trials.

| SOA (ms) | Predictor | | | $F$-value | df<sub>1</sub>, df<sub>2</sub> | $p$-value | $\eta_p^2$ |
| --- | --- | --- | --- | --- | --- | --- | --- |
| 0 | Bilateral Pre-stimulus Alpha Power |  |  | 0.38 | 2, 161 | 0.685 | 0.005 |
|  | Pre-stimulus Alpha Lateralization |  |  | 0.45 | 2, 162 | 0.636 | 0.006 |
|  | Ipsilateral Pre-stimulus Alpha Power |  |  | 2.02 | 2, 162 | 0.136 | 0.024 |
|  | Contralateral Pre-stimulus Alpha Power |  |  | 2.01 | 2, 162 | 0.138 | 0.024 |
| 120 | Bilateral Pre-stimulus Alpha Power |  |  | 3.25 | 2, 160 | 0.042<sup>*</sup> | 0.039 |
|  | Pre-stimulus Alpha Lateralization |  |  | 1.89 | 2, 161 | 0.154 | 0.023 |
|  | Ipsilateral Pre-stimulus Alpha Power |  |  | 3.97 | 2, 161 | 0.021<sup>*</sup> | 0.047 |
|  | Contralateral Pre-stimulus Alpha Power |  |  | 0.87 | 2, 161 | 0.421 | 0.011 |
| 1240 | Bilateral Pre-stimulus Alpha Power |  |  | 0.007 | 2, 161 | 0.993 | 0.001 |
|  | Pre-stimulus Alpha Lateralization |  |  | 0.64 | 2, 160 | 0.531 | 0.008 |
|  | Ipsilateral Pre-stimulus Alpha Power |  |  | 1.04 | 2, 160 | 0.355 | 0.013 |
|  | Contralateral Pre-stimulus Alpha Power |  |  | 0.17 | 2, 159 | 0.847 | 0.002 |
| Post-hoc contrasts |  |  |  |  |  |  |  |
| SOA (ms) | Predictor | Level 1 | Level 2 | $F$-value | df<sub>1</sub>, df<sub>2</sub> | $p$-value | Cohen's $d$ |
| 120 | Bilateral Alpha | Weak | Medium | 5.07 | 1, 160 | 0.026<sup>*</sup> | +0.43 |
|  | Bilateral Alpha | Weak | Strong | 4.67 | 1, 160 | 0.032<sup>*</sup> | +0.42 |
|  | Bilateral Alpha | Medium | Strong | 0.009 | 1, 160 | 0.925 | +0.02 |
|  | Ipsilateral Alpha | Weak | Medium | 0.27 | 1, 161 | 0.601 | −0.10 |
|  | Ipsilateral Alpha | Weak | Strong | 4.51 | 1, 161 | 0.035<sup>*</sup> | +0.41 |
|  | Ipsilateral Alpha | Medium | Strong | 7.09 | 1, 161 | 0.009<sup>**</sup> | +0.51 |

Top: Results of one-way LMEs predicting $d'$ from each alpha power predictor (tertile bins) within neutral pre-cue trials only. Bottom: Post-hoc pairwise contrasts for significant effects at 120 ms. Significant effects are indicated with asterisks (\* = $p < 0.05$ ; \*\* = $p < 0.01$ ).

### Aperiodic activity

Aperiodic offset and exponent did not differ between correct and incorrect trials at any of the six bilateral alpha-binning channels (all *p ≥* 0.32, see Table S1), indicating that the alpha power effects reported above are not associated with broadband shifts in aperiodic activity.

### Gaze analysis

Our results showed that participants responded more accurately at the 120 ms target-to-post-cue SOA when ipsilateral pre-stimulus alpha relative to the target location was increased. As alpha lateralization is associated with eye movements and gaze shifts, we tested whether small gaze shifts before target onset were associated with increased performance when the gaze shift was in the direction of the upcoming target (Mössing et al., 2024; Popov et al., 2023; Van Ede et al., 2019). We tested this for each pre-cueing condition separately for targets presented in the left and right hemifield. A descriptive inspection of the gaze density heatmaps showed no systematic difference in the spatial distribution of gaze density between correct and incorrect trials at the 120 ms target-to-post-cue SOA for the neutral and invalid precueing condition (see Figure S1C–F). No pixel survived FDR correction for the difference in gaze density between correct and incorrect trials in the neutral and invalid pre-cueing condition (all *q ≥* 0.137; see Table S2). This indicates that the difference in pre-stimulus gaze position between correct and incorrect trials cannot account for the effects of pre-stimulus alpha power in the neutral and invalid pre-cueing conditions reported above. There was a significant difference in gaze density in the valid pre-cueing condition, with stronger gaze density in the center of the screen being associated with worse performance (see Figure S1A–B and Table S2). However, as there were no systematic biases towards the direction of the upcoming target and as the valid pre-cueing condition is not particularly relevant for our pre-stimulus alpha analysis we treat this result as incidental.

## Discussion

Iconic memory—the earliest stage of visual memory—is modulated both by selective spatial attention (P. J. C. Smith & Busch, 2026) and by fluctuations in spontaneous pre-stimulus brain states (P. J. C. Smith & Busch, 2025b). This study tested whether the effects of attentional cueing and spontaneous pre-stimulus brain states reflect a shared inhibitory process or independent contributions. To this end, we combined an endogenous spatial pre-cue indicating the hemifield of the upcoming stimulus display with a typical iconic memory paradigm, where a partial-report post-cue indicated the to-be-reported target stimulus. Our analysis focused on the effects of spatial cueing and alpha-band lateralization on performance at the 120 ms target-to-post-cue SOA. This SOA best reflects a time point at which the stimulus has already disappeared, but stimulus information is still available in iconic memory, and performance has not yet settled at its short-term memory asymptote. This makes it the window that most cleanly indexes the decaying iconic trace. Here, both invalid attentional pre-cues and weaker ipsilateral pre-stimulus alpha power impaired performance. These two effects were independent and additive rather than interactive.

The behavioral data reproduced the central finding of P. J. C. Smith and Busch (2026). Pre-cue validity modulated *d^′^* at the earliest target-to-post-cue SOAs, and this modulation was carried entirely by a cost at invalidly cued locations relative to the neutral, distributed-attention baseline, with no detectable benefit at validly cued locations. As expected, pre-cue validity also produced conventional cue-driven alpha lateralization, with greater ipsi- and lower contralateral alpha power relative to the attended hemifield (see Figure 1B).

The pre-stimulus alpha power results reveal an inhibitory pattern. At 120 ms, stronger pre-stimulus alpha power ipsilateral to the target predicted higher *d^′^*. By contrast, contralateral pre-stimulus alpha power relative to the target had no effect, at any target-to-post-cue SOA (Figure 2B). Restricting the analysis to neutral trials reproduced the same selective ipsilateral effect. Directed spatial attention is absent in these trials; therefore, this closely replicates the uncued, spontaneous lateralization effect of P. J. C. Smith and Busch (2025b). The selective ipsilateral effect likely reflects suppression of the cortical hemisphere representing the irrelevant hemifield, rather than facilitation of the hemisphere representing the target. Similar lateralization patterns have been observed in response to attentional cues (Foxe & Snyder, 2011; Händel et al., 2011; Jensen & Mazaheri, 2010; Thut et al., 2006; Worden et al., 2000), and spontaneous trial-by-trial lateralization has been shown to predict endogenous shifts of spatial attention (Balestrieri & Busch, 2022; Bengson et al., 2014; Nadra et al., 2023). This links the spontaneous and cue-driven effects to a common inhibitory pattern.

This ipsilateral effect did not interact with pre-cue validity; instead, both factors contributed additive main effects at 120 ms. If cue-induced and spontaneous alpha lateralization were a single mechanism, the benefit of ipsilateral alpha should have depended on cue validity, for instance increasing under valid pre-cues and decreasing under invalid pre-cues. The absence of such an interaction points to two partly independent inhibitory routes to the same endpoint: endogenous, cue-driven allocation of spatial attention, and ongoing fluctuations in cortical excitability indexed by spontaneous alpha lateralization. Both bias processing toward the relevant hemifield by inhibiting the irrelevant one. These effects could reflect endogenous attention—one cued and the other spontaneously self-initiated with both shaping iconic memory (Nadra et al., 2023). Alternatively, the spontaneous effect may not be exclusive to deliberate attention, but instead reflect a more general property of the momentary excitability state of the early visual cortex, to which directed attention is one contributor (Iemi et al., 2022).

This influence of alpha power relates to the neural basis of iconic memory. Iconic memory depends on the persistent firing of neurons in early visual cortex (Teeuwen et al., 2021), and the gain of these responses is inversely related to ongoing alpha power (Dougherty et al., 2017; Iemi et al., 2019). Lateralized alpha power is therefore well placed to determine, hemifield by hemifield, which sensory information persists long enough to be reported. On this account, strong alpha power ipsilateral to the target suppresses processing of the irrelevant hemifield and protects the target trace, improving performance; an invalid cue drives the same inhibitory state toward the wrong hemifield, suppressing the target trace and producing the behavioral cost. The cost of an invalid cue and the advantage conferred by strong ipsilateral alpha power may thus be two expressions of the same inhibitory mechanism. This suppression-over-enhancement pattern is consistent with accounts of noise exclusion in spatial attention, where performance is limited primarily by the observer’s ability to filter out irrelevant information rather than by the quality of the target representation itself (Dosher & Lu, 2000). The same logic plausibly extends to the absence of any facilitatory effect of contralateral alpha power, since excitability at the processing hemisphere is not the bottleneck if suppression of the irrelevant hemifield is what matters.

The effect of ipsilateral pre-stimulus alpha power relative to the target at the 0 ms target-to-postcue SOA was less straightforward. Here, intermediate ipsilateral pre-stimulus alpha power relative to the target was associated with significantly lower *d^′^* than both weak and strong power, which did not differ from one another. This non-monotonic pattern does not follow the graded inhibitory relationship observed at 120 ms. However, the lateralization model at 0 ms told a more orderly story: relatively stronger ipsilateral than contralateral pre-stimulus alpha power was associated with better performance, consistent with suppression of the irrelevant hemifield. Because the lateralization index reflects the relative balance between ipsilateral and contralateral power rather than ipsilateral power alone, the two results are not in conflict: a given hemifield’s relative advantage can be graded and orderly even when the absolute level of ipsilateral power, considered on its own, is not.

At the latest target-to-post-cue SOA of 1240 ms, the effect of pre-stimulus lateralization reversed: relatively stronger contralateral than ipsilateral pre-stimulus alpha power was associated with better performance. This may mark the handover from iconic memory to the consolidation of information into visual short-term memory, a later stage at which attentional facilitation, rather than inhibition, becomes the more relevant mechanism (Botta et al., 2019; Myers et al., 2017; Souza & Oberauer, 2016). This is similar to the behavioral results of our previous work, which shows that the attentional cost associated with invalid compared to neutral pre-cues is no longer present at the target-to-post-cue SOA of 1240 ms (P. J. C. Smith & Busch, 2026).

These results bear on the long-standing question of whether iconic memory—the earliest stage of visual memory—is pre-attentive. Accounts grounded in the distinction between phenomenal and access consciousness hold that iconic memory reflects phenomenally rich, pre-attentive contents, with attention entering only at the transfer to short-term memory or at an intermediate fragile store (Block, 2011; Lamme, 2004, 2006; Pinto et al., 2013; Sligte et al., 2008). Our findings rule out a strictly pre-attentive account: both an explicit attentional manipulation and an endogenous neural index of spatial selection modulated performance at early target-to-post-cue SOAs, i.e., the onset of iconic memory itself. The iconic memory trace is impaired by attentional allocation towards the irrelevant hemifield—favoring a prominent role of attention in iconic memory formation. These results are consistent with recent behavioral work showing that manipulations of spatial attention impact iconic memory performance (Botta et al., 2019; Botta et al., 2023; P. J. C. Smith & Busch, 2026). Iconic memory is genuinely sensitive to spatial attention, and this sensitivity takes the form of inhibitory selection against irrelevant locations.

## Conclusion

By combining endogenous spatial cueing with pre-stimulus alpha analysis in a single partial-report paradigm, we consolidate two previously separate observations—a behavioral attentional cost (P. J. C. Smith & Busch, 2026) and a spontaneous alpha-lateralization effect (P. J. C. Smith & Busch, 2025b)— within one account of how spatial selection shapes the earliest stage of visual memory. Both an invalid attentional cue and weak ipsilateral pre-stimulus alpha relative to the target impaired performance at the 120 ms target-to-post-cue SOA, and both are consistent with suppression of irrelevant locations. That the two effects were additive rather than interactive indicates that both cue-directed attention and ongoing cortical excitability contribute to this inhibitory shaping. Iconic memory is thus sensitive to attention and continuously biased by the inhibitory state of early visual cortex.

## Data and code availability

Data will be available at the Open Science Framework (https://osf.io/b95zr). Code will be available at https://github.com/pauljcs/alpha-attention-iconic.

## Acknowledgments

This work was supported by a grant from the German Research Foundation (DFG; BU 2400/15-1). We thank Isabel Amelie Finken, Bernhard Matthias Meier, Maya Müllers, Mathilde Maria Pöppelmann, Philipp Konrad Rapp, Felix Rinn, and Simon Steibel for help with the data acquisition.

## Supplementary Material

**Figure S1:**
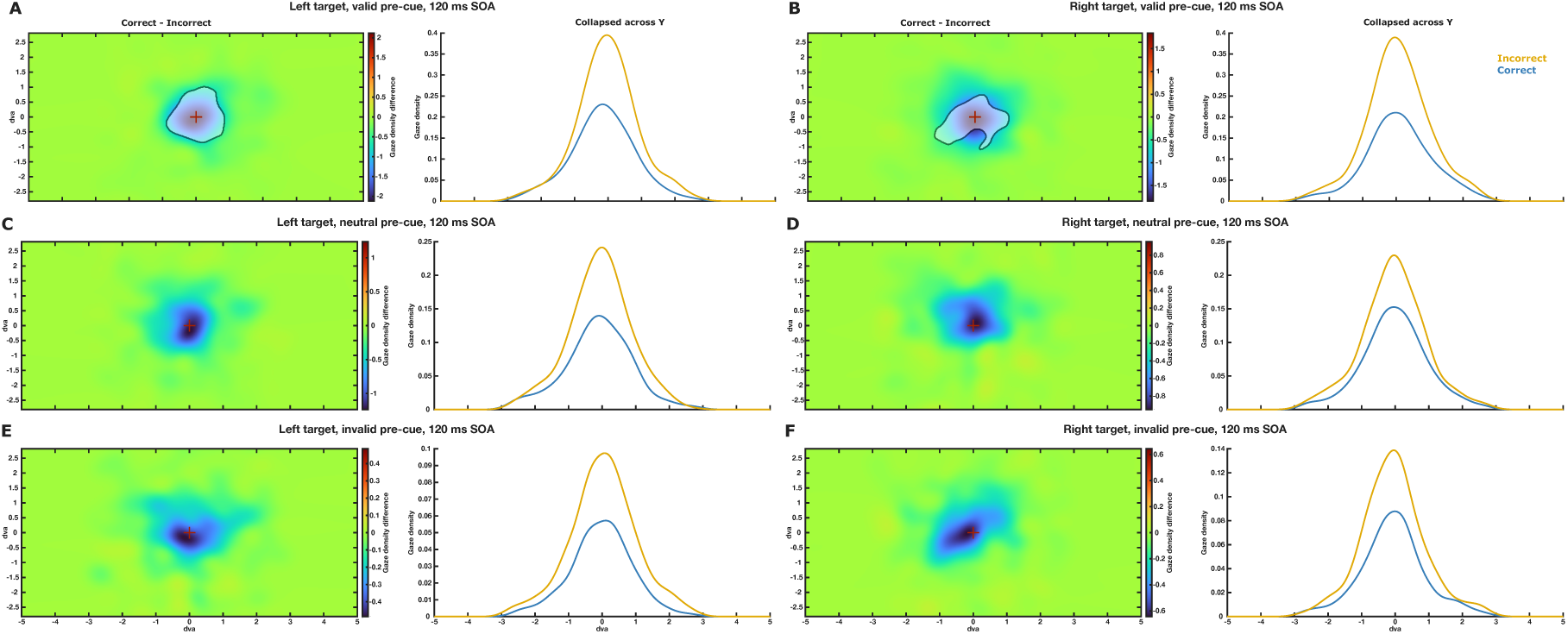
**A**: Gaze density for correct vs. error responses for valid pre-cue trials with targets presented in the left hemifield at the 120 ms SOA. Left plot shows a heatmap of the gaze density difference for correct vs. error responses. Warm values indicate stronger gaze density for correct trials and cold colors indicate stronger gaze density for incorrect trials. Right plot shows horizontal gaze position separately for correct vs. error responses. Plot area is restricted to the center of the screen. The center of the screen is indicated via a red fixation cross. Pixel regions showing significant differences are indicated with a grey mask. The following panels follow these conventions. **B**: Gaze density for correct vs. error responses for valid pre-cue trials with targets presented in the right hemifield at the 120 ms SOA. **C**: Gaze density for correct vs. error responses for neutral pre-cue trials with targets presented in the left hemifield at the 120 ms SOA. **D**: Gaze density for correct vs. error responses for neutral pre-cue trials with targets presented in the right hemifield at the 120 ms SOA. **E**: Gaze density for correct vs. error responses for invalid pre-cue trials with targets presented in the left hemifield at the 120 ms SOA. **F**: Gaze density for correct vs. error responses for invalid pre-cue trials with targets presented in the right hemifield at the 120 ms SOA.

**Table S1:** Aperiodic offset and exponent, correct vs. incorrect trials, by channel.

| Channel | Parameter | $t$ | $p$ |
| --- | --- | --- | --- |
| 18 | Offset | -0.82 | 0.418 |
|  | Exponent | 0.20 | 0.842 |
| 19 | Offset | -0.66 | 0.516 |
|  | Exponent | 0.14 | 0.889 |
| 22 | Offset | -0.33 | 0.746 |
|  | Exponent | 0.30 | 0.767 |
| 26 | Offset | -0.64 | 0.525 |
|  | Exponent | 0.44 | 0.664 |
| 28 | Offset | 0.99 | 0.328 |
|  | Exponent | 0.91 | 0.369 |
| 57 | Offset | 0.07 | 0.943 |
|  | Exponent | 0.58 | 0.563 |

Paired two-tailed $t$ -tests comparing aperiodic offset and exponent between correct and incorrect trials at each of the six bilateral alpha-binning electrodes (P3, P5, P6, P8, PO3, and PO8), in the pre-stimulus window (-1000:-2 ms before stimulus onset).

**Table S2:** Gaze density comparison (correct vs. incorrect trials) at the 120 ms SOA.

| Cue | Target hemifield | Peak $\vert t\vert $ | Min. $p_{\text{unc}}$ | Min. $q_{\text{FDR}}$ |
| --- | --- | --- | --- | --- |
| Valid | Left | 6.03 | $< .001$ | $< .001$ |
| | Right | 5.27 | $< .001$ | 0.011 |
| Neutral | Left | 4.91 | $< .001$ | 0.137 |
| | Right | 4.71 | $< .001$ | 0.249 |
| Invalid | Left | 3.99 | $< .001$ | 0.687 |
| | Right | 4.37 | $< .001$ | 0.725 |

Per-pixel two-sided $t$ -tests comparing two-dimensional gaze density between correct and incorrect trials in the pre-stimulus window (-1000 to -2 ms), corrected across pixels with the false discovery rate (FDR; $q < 0.05$ ). Peak $|t|$ is the largest absolute $t$ across pixels; Min. $p_{\text{unc}}$ and Min. $q_{\text{FDR}}$ are the smallest uncorrected and FDR-corrected values, respectively.

